# Optical recording of action potential initiation and propagation in mouse skeletal muscle fibers

**DOI:** 10.1101/292391

**Authors:** Q. Banks, S.J.P. Pratt, S.R. Iyer, R.M. Lovering, E.O. Hernández-Ochoa, M.F. Schneider

## Abstract

Individual skeletal muscle fibers have been used to examine a wide variety of cellular functions and pathologies. Among other parameters, skeletal muscle action potential propagation has been measured to assess the integrity and function of skeletal muscle. In this paper, we utilize Di-8-ANEPPS, a potentiometric dye and mag-fluo-4, a low-affinity intracellular calcium indicator to non-invasively and reliably measure action potential conduction velocity in skeletal muscle. We used an extracellular bipolar electrode to generate an electric field that will initiate an action potential at one end of the fiber or the other. Using enzymatically dissociated flexor digitorum brevis (FDB) fibers, we demonstrate the strength and applicability of this technique. Using high-speed line scans, we estimate the conduction velocity to be approximately 0.4 m/s. In addition to measuring the conduction velocity, we can also measure the passive electrotonic potentials elicited by pulses by either applying tetrodotoxin (TTX) or reducing the bath sodium levels. We applied these methodologies to FDB fibers under elevated extracellular potassium conditions, and found that the conduction velocity is significantly reduced compared to our control concentration. Lastly, we have constructed a circuit model of a skeletal muscle in order to predict passive polarization of the fiber by the field stimuli. Our predictions from the model fiber closely resemble the recordings acquired from *in vitro* assays. With these techniques, we can examine how many different pathologies and mutations affect skeletal muscle action potential propagation. Our work demonstrates the utility of using Di-8-ANEPPS or mag-fluo-4 to non-invasively measure action potential conduction velocity.

## INTRODUCTION

The conduction velocity of the muscle fiber action potential (AP) is a valuable parameter in physiology. It reflects the integrity and excitability of the sarcolemma and transverse tubules and could show alterations in different pathologies [1] [2]. If a normal muscle fiber is briefly depolarized over a local area, an AP will be initiated at the point of depolarization and will propagate longitudinally along the sarcolemma, and radially throughout the T-tubule network. Conduction velocity could be affected by disruption or dysregulation of the T-tubule network [3], ion channels and other proteins [4], among numerous possibilities. If the depolarization or repolarization phase of the muscle AP is affected, it is possible that conduction velocity has also been affected. Numerous studies have already looked at conduction velocity of muscle, and how it changes under different conditions and pathologies [5] [6] [7]. Past studies examining conduction velocity have used intracellular microelectrodes techniques, or electromyography (EMG) in living animals. These methods have allowed us to understand many different aspects of cellular physiology, but they have their pitfalls. Microelectrode impalement or patch clamp pipette in the whole cell configuration could disrupts the sarcolemma and intracelluar milleu. EMG records from gross muscle, are ideal for the study of muscle fiber groups, but not for studies in in individual cells.

Previous research in our lab has shown that some fibers display a local twitch response at alternating ends of the fiber when stimulating with alternating polarity using remote bipolar electrodes [8]. Stimulation at one polarity caused a local contraction response only at the end of the fiber facing the negative electrode, while switching the polarity caused a local contraction at the other end of the fiber. After dissociation, some fibers display this alternating response to alternating polarity stimuli. We are also able to induce the alternating end local contractile response to alternating polarity field stimuli by either applying tetrodotoxin (TTX) or removing Na^+^ [8] From this, we deduced that negative polarity field stimulation initiated a local depolarization, resulting in local release of Ca2+, at the end of the muscle fiber facing the negative electrode, while the other polarity elicited a local hyperpolarization and thus no contraction, consistent with passive opposite polarity local electronic polarization at the ends of the fibers. The above observations, allowed us to hypothesize that in fully excitable fibers, the electric field generated by bipolar stimulation initiates an AP at the end of the fiber facing the negative electrode (Figure 1a). Application of voltage to the field electrodes, will initially causes a passive hyperpolarization at the end of the fiber near the positive electrode. Likewise, the end closer to the negative electrode undergoes a passive depolarization. If sufficierntly large, this depolarization will trigger an AP, which will propagate in the opposite direction than the field-induced current.

**Figure 1.**
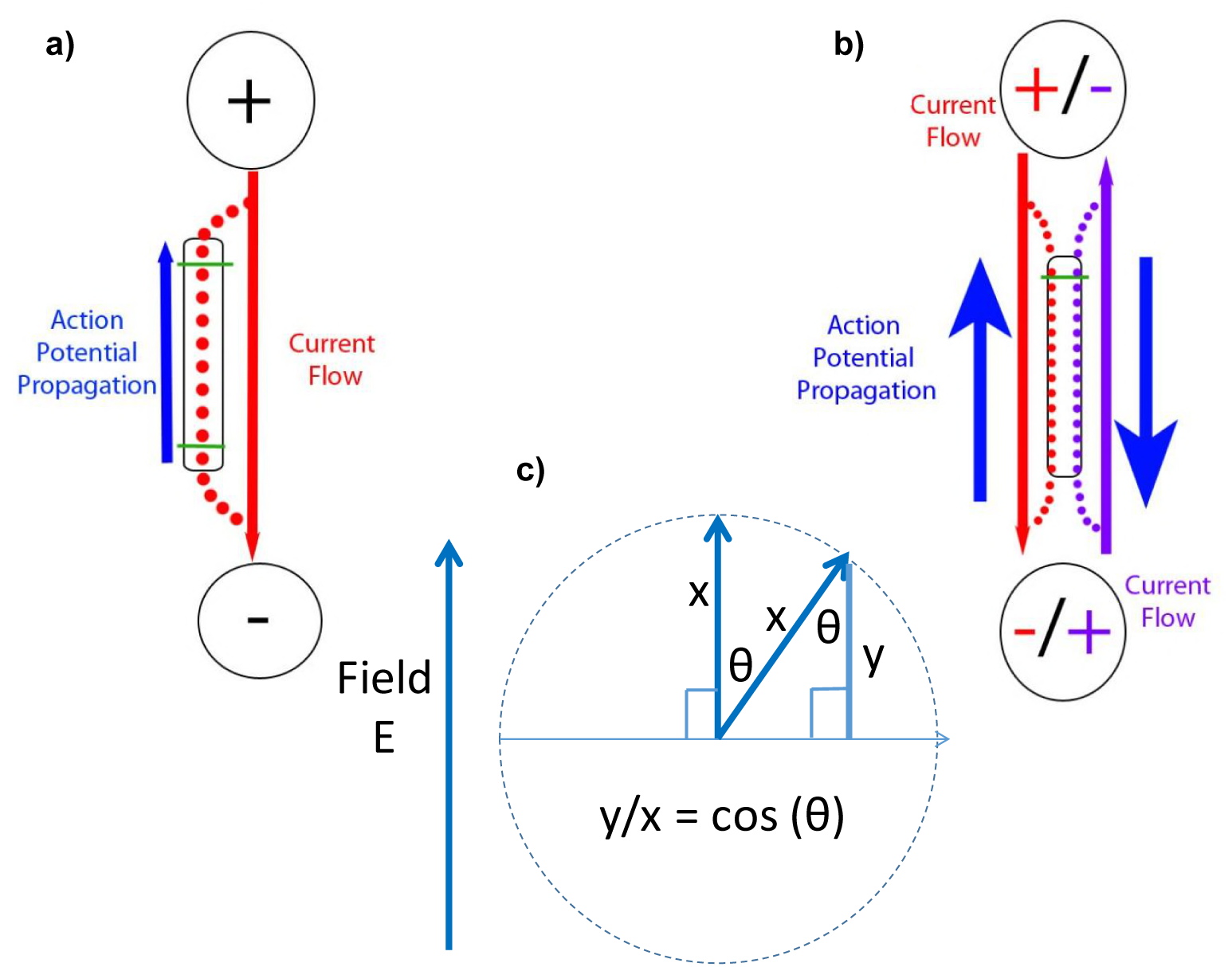
Effects of electric field generation on isolated muscle fibers. a) Upon stimulation from a pair of external remote bipolar electrodes, the two ends of the muscle fiber will be affected in opposite ways. The end near the positive pole will be hyperpolarized (made more negative; V_m_ is measured by internal voltage relative to external voltage), while the end near the negative pole will be depolarized, and this is where the action potential will be initiated. We can change our recording location from one end to the other end, thereby examining the initiation and propagation of the action potential. b) Alternatively, by changing the polarity of the electrodes, we can cause alternate depolarization of the fibers ends, hence the location of the action potential initiation, while recording at the same fiber end, allowing for the measurement of propagation velocity of action potentials or action potential induced-Ca2+ transients, providing that the fiber length is known. c) A vector diagram showing the effect of the electric field on muscle fibers when they are not parallel to each other. If within 45 degrees of the field, fibers will experience at least cos(45) = 0.71 of the applied field.

Here we take advantage of this phenomenon to measure skeletal muscle AP propagation. We used the membrane impermeable, potentiometric dye, 1-(3-sulfonatopropyl)-4[β[2-(Di-*n*-octylamino)-6-naphtyl]vinyl]pyridinium betaine (Di-8-ANEPPS) to monitor changes in membrane potential in response to field stimulation. The rationale is as follows. If we optically measure the AP using high speed resolution (i.e., high-speed line scans at both ends of the muscle fiber) we can record the initiation (at the negaive facing end of the fiber) and the propagated AP (at the positive facing end of the fiber). Alternatively, we can switch the polarity of the bipolar electrodes, and record line scans from the same fiber end for both negative and positive stimuli (i.e., for both depolarizing and hyperpolarizing field stimuli; Figure 1b).

If a muscle fiber is not oriented parallel to the electric field it will only experience a fraction of the full field along its length. Consider a muscle fiber of length X oriented parallel to an electric field E. According to Fig 1C, where X is the radius of a circle, if the fiber is oriented at an angle θ to the applied field, the fiber would experience an effective fractional component of the field (y/x) equal to cos(θ) along its length, which is the effective longitudinal field for the fiber at angle θ to the field. In our experiments we only record from fibers with θ < 45 degrees, in which case the fibers experience at least cos(45) = 0.71 of the applied field.

We can also use Ca^2+^ dyes, like mag-fluo-4 to estimate the propagated AP velocity. APs are soon followed by calcium release from the SR, and the resulting calcium transients also propagate towards the end of the fiber being hyperpolarized.

Fluorescent dyes can target individual cells, and when used at low concentrations, apperar to not disrupt the standard functionality of cells [9]. Fluorescent measures have already been used to examine many different aspects of cellular structures and functions [10–16]. We have refined a method to non-invasively examine and estimate the conduction velocity in isolated skeletal muscle fibers. In this paper, we demonstrate the functionality and viability of this method, by challenging flexor digitorum brevis (FDB) fibers to elevated extracellular potassium.

Elevated extracellular potassium can arise, among other mechanisms, from extensive muscle use and fatigue, depolarizing the membrane, reducing the amount of Ca2+ released, and diminishing the level of tension that can be achieved [17, 18]. Altering the ionic composition on either side of the membrane will affect the driving forces for the ions, so it stands to reason that the conductance of ions and the conduction velocity will be affected as well. Here we test the effect of elevated extracellular potassium on conduction velocity under field stimulation conditions, and find that it is reduced in a time dependent manner, and that a number of fibers become too depolarized to initiate an AP. The above observations demonstrate the utility of this new method to estimate AP conduction in a non-invasive manner in single muscle fibers. Part of this work was presented at the Biophysical Society Meeting, 2018.

## MATERIALS AND METHODS

### Ethical approval

All animals were housed in a pathogen-free area at the University of Maryland, Baltimore. The animals were euthanized according to authorized procedures of the Institutional Animal Care and Use Committee, University of Maryland, Baltimore, by regulated delivery of compressed CO_2_ overdose followed by cervical dislocation.

### FDB Fiber Preparation

FDB muscles were isolated from the hindlimbs of female CD1 mice between 4 and 8 weeks old. A total of 24 mice were used. Fibers were placed in 2 mg/mL type I collagenase/minimum essential medium (MEM) (Gibco, Carlsbad, CA, Cat. No. 11095098) to enzymatically dissociate single fibers for 4 hours. Afterward, FDBs were also dissociated via mechanical means using a glass pipette in MEM with 0.1% gentamycin (Sigma, St. Louis, MO; Cat. No. G1397) and 10% fetal bovine serum (Gemini Bio-Products, West Sacramento, CA, Cat. No. 100-106). Single fibers were plated on laminin-coated glass bottom dishes (Matek Cor. Ashland, MA, Cat. No. P35G-1.0-14-C) in MEM with 0.1% gentamycin. Dishes were placed in a 37^°^ C with 5% CO_2_ incubator overnight. Cells were tested within 48 hours of isolation.

### T-Tubule Network Imaging

Isolated cells were stained with 5 µM of Di-8-ANEPPS in MEM for 2 hours. Cells were kept in the incubator during this time. Excitation for Di-8-ANEPPS was provided by a 543 nm laser, and emitted light was collected at > 560 nm. The laser intensity was set to 20%. Cells were examined for structural integrity before being tested. Fibers were imaged using a 60x water-immersion objective lens with MEM as the recording solution. A set of four scans were taken at 3.5x zoom and averaged to produce each frame image. Images were background corrected to better display the T-tubule network.

### Di-8-ANEPPS Voltage Signal Recording

Isolated cells were stained with 5 µM of Di-8-ANEPPS (Invitrogen, Carlsbad, CA, Cat No. D3167) in MEM for 2 hours. Cells were kept in the incubator during this time. MEM was aspirated and replaced with L-15 directly before testing. Cells were stimulated at 12 volts for 0.5 ms while line scans were taken at rates of either 10,000 lines/sec or 50,000 lines/sec (512 x 10,000 or 50,000 pixels). Records are an average of 6 scans from the same fiber at a given location. TTX (Millipore Corp., Darmstadt, Germany, Cat. No. 554412) was administered at a 1 µM concentration. TTX was allowed 10 minutes after application to block Na^+^ channels and obtain passive responses to stimulation. Excitation for Di-8-ANEPPS was provided by a 532 nm laser, and emitted light was collected at >550 nm. To improve signal/noise, excitation light intensity was set to 20% of the 532 nm laser in the confocal system. To minimize photo damage, excitation light exposure was limited to 30 ms, beginning 3 ms prior to the start of the field stimulus. Cells were examined for structural integrity and twitch responses to field stimulation before being tested. Fibers were imaged using either a 10x or 60x water-immersion objective lens with L-15 as the recording solution. Images were background corrected by subtracting an average value recorded outside the cell. The average fluorescence before stimulation was used to find F_0_ (baseline before stimulation), which was used to scale the Di-8-ANEPPS signal in the same ROI to obtain ΔF/F0 (change in fluorescence compared to baseline). Experiments were performed at room temperature, 21-23 C°.

### Mag-Fluo-4 Ca2+ recordings

Isolated cells were stained with 1 µM of mag-fluo-4 (Life Technologies, Eugene, OR, Cat. No. M14206) in 37o Co L-15 (Gibco, Carlsbad, CA, Cat. No. 21083027) with 0.1% gentamycin and 0.25% albumin (Sigma-Aldrich, St. Louis, MO, Cat No. A8806-5G) for 30 minutes. Media was aspirated and replaced with standard room temperature L-15 for 30 minutes before testing. Cells were stimulated at 15 volts for 0.5 while line-scans (*xt)* were taken at a rate of 10,000 lines/sec (512 × 10,000 pixels). Excitation for mag-fluo-4 was provided by a 488-nm laser. The emitted light was collected at >505 nm. Cells were examined for structural integrity and twitch response to field stimulation before being tested for conduction velocity. Fibers were imaged using either a 10× or 63× water-immersion objective lens with L-15 as the recording solution. Images were background corrected by subtracting an average value recorded outside the cell (Figure 2a). The average fluorescence before stimulation was used to find F_0_, which was used to scale the mag-fluo-4 signal in the same ROI to obtain ΔF/F_0_. Experiments were performed at room temperature, 21-23 C×.

**Figure 2.**
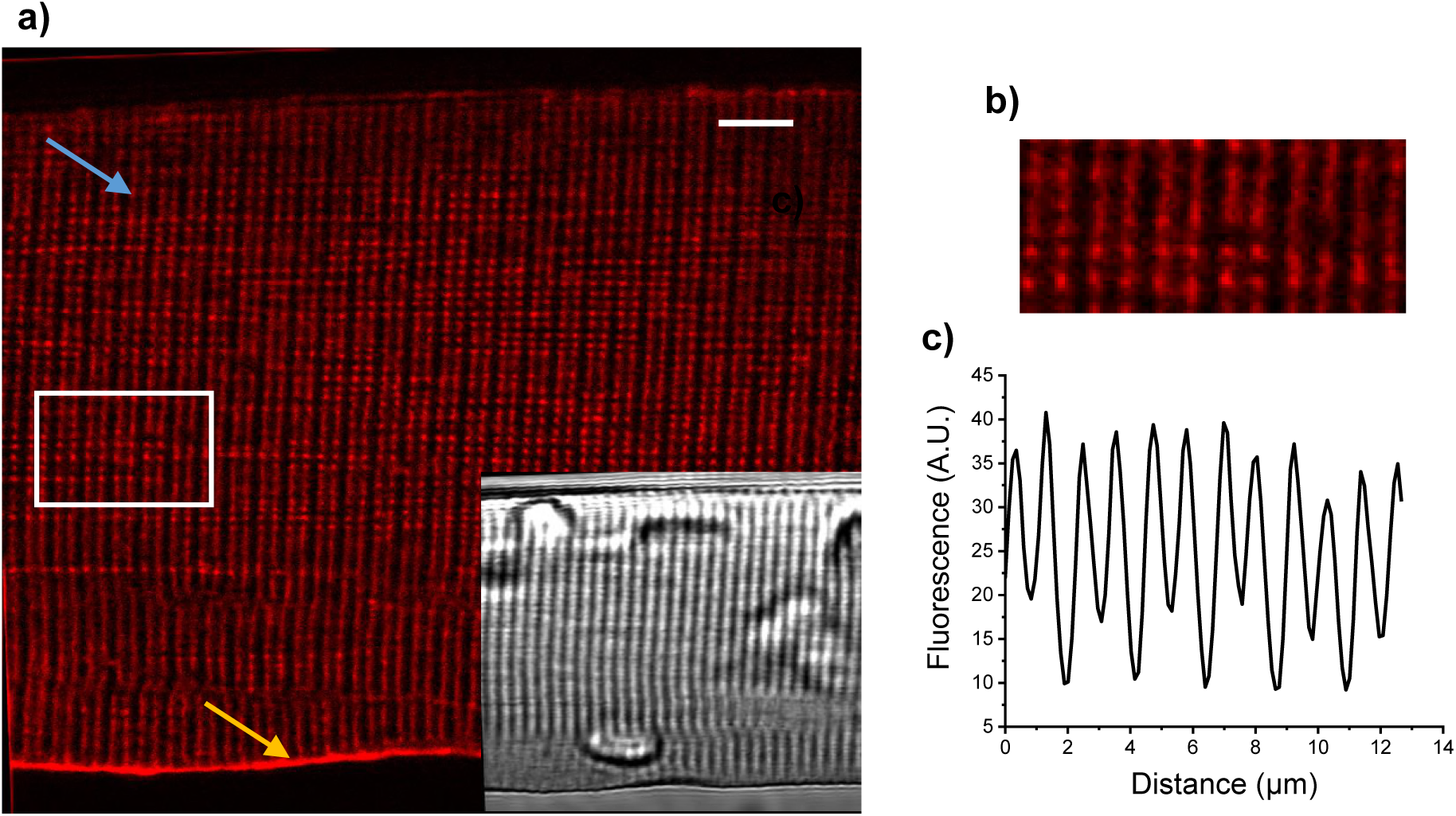
Di-8-ANEPPS staining in a single skeletal muscle fiber. a) Representative confocal image of a segment of a FDB fiber stained with Di-8-ANEPPS to visualize the T-tubule system. Calibration bar is 5 μm. Transmitted light is shown in the inset. Arrows indicate the location of regions of interest that could potentially be used to measure the time course of surface action potentials (orange arrow) or core action potentials (blue arrow) used in Fig. 3. b) Zoomed in segment of image in panel a) indicated by white box. c) Graph indicates average fluorescence intensity of boxed region shown in b). The space between T-tubules is approximately 2 μm. Distance between the troughs represents the distance from Z line to Z line.

### Data Analysis

Conduction velocity data were acquired in LSM 5 Live (Zeiss, Jena, Germany), processed in Excel (Microsoft, Redmond, WA, USA), and analyzed and plotted using OriginPro 2016 (OriginLab Corporation, Northampton, MA, USA). Data following a normal distribution are presented as box plots indicating the range, Q1, mean, Q3 (solid lines), and median (smaller box). Where noted, a two-sample t-test or paired-sample t-test was used to test for significance (set at *p* < 0.05). Normality was evaluated using the Shapiro-Wilk test (p < 0.05: reject normality). For samples that were had normality rejected, a Mann-Whitney test was performed (set at p < 0.05). Non-parametric data are represented as box plots indicating the range, Q1, Q3, (solid lines), and median (smaller box). T-tubule network images were acquired using FluoView (Olympus, Center Valley, PA, USA). Images were processed using ImageJ (National Institutes of Health, Rockville, MD).

## RESULTS

### 2-D Visualization of T-Tubule Network

Proper AP propagation requires a uniform, structurally intact T-tubule network, as the AP propagates radially through the T-tubules as well as along the sarcolemma. Di-8-ANEPPS binds to the cell membrane and diffuses into the T system and binds to the T-Tubule membrane. Thus it can be used to measured transmembrane potentials of both the surface membrane and T-Tubule network. Figure 2a shows a *xy* image of the T-Tubule network as shown via Di-8-ANEPPS. At a 5 µM concentration of Di-8-ANEPPS, the transverse tubules, as well as the longitudinal tubules, were well defined. Figure 2b shows a close up of a small section of the T-tubule network, and the normal spacing between tubules. Figure 2c shows the fluorescence intensity profile of Figure 2b.

### Passive and Active Responses to brief (0.5 ms) alternate polarity field stimulation

When using field stimulation to elicit an AP, we will produce both passive and active electrical responses. The passive responses arise from electrotonic polarization of the membrane capacitance and resistance of the resting membrane, and the active responses encompass the AP resulting from the electrotonic depolarization. Using Di-8-ANEPPS (Figure 3a), we can optically record both types of membrane responses over time (Figure 3b). Signals from Di-8-ANEPPS are small, so we averaged a total of 6 recordings of each polarity at each acquired location on the fiber to improve signal/noise. A total of 30 fibers from 13 mice were included in this section of the study. As mentioned earlier, when recording from the ends of a fiber, stimulation will cause AP [4] initiation at the end of the fiber facing the negative field electrode, and the AP will conduct to the opposite end. When recording from the middle, however, the conduction delay is the same from both ends of the fiber and there is no electronic polarization prior to the AP. Figure 3b displays example averaged recordings from both ends and the middle of the fiber for each polarity of field stimulation. TTX was applied after active responses were acquired in order to block AP initiation or propagation and isolate passive hyperpolarizations and depolarizations. Example records are displayed in Figure 3c. Previous studies have performed Di-8-ANEPPS calibration tests using voltage steps [19, 20]. Both studies used a holding potential of −90 mV. Based on their studies, they estimated that the amplitude of APs observed through Di-8-ANEPPS is approximately between 112 mV and 123 mV. This set of experiments showed that we can reliably examine changes in membrane voltage with Di-8-ANEPPS.

**Figure 3.**
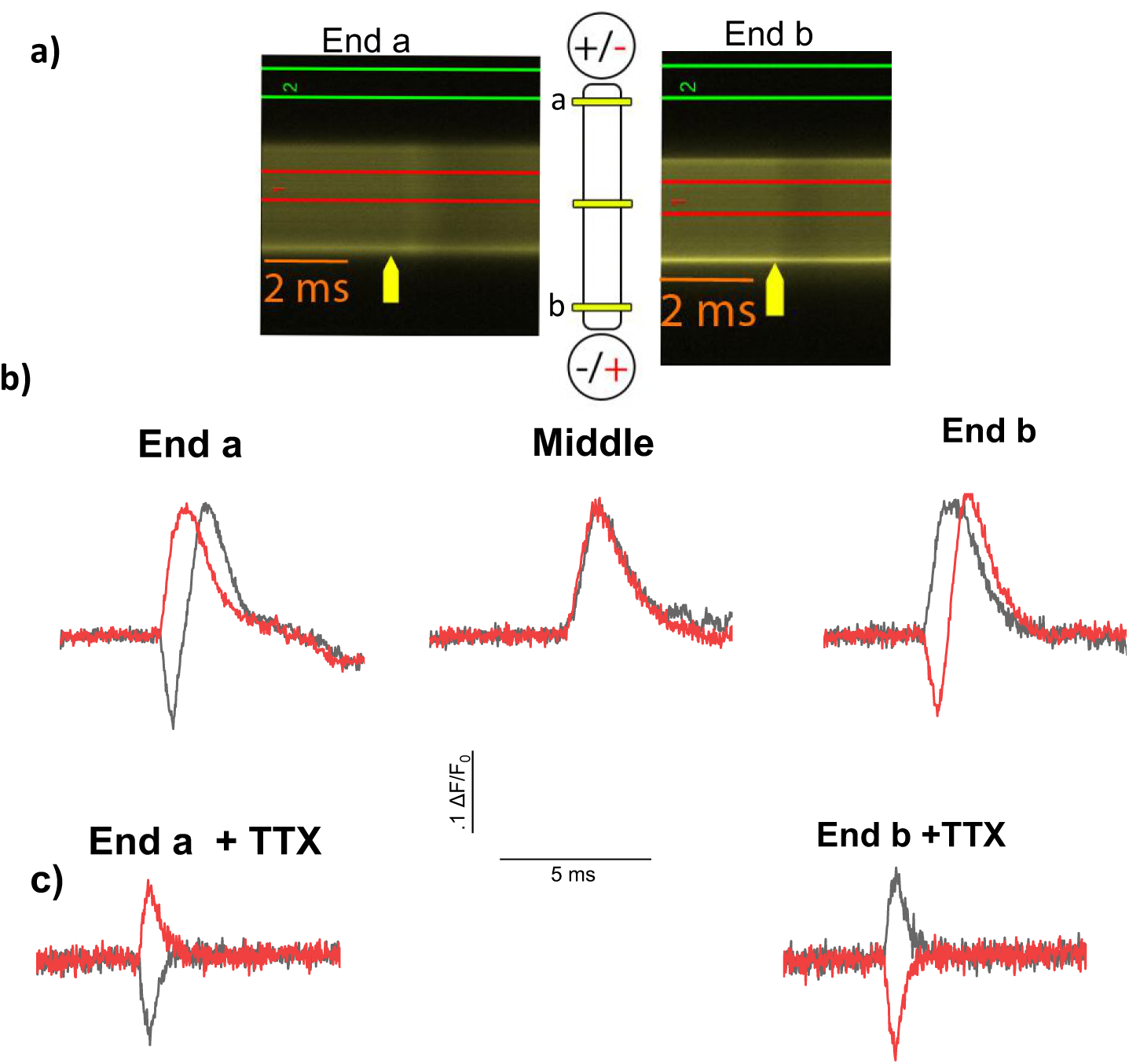
Averaged (n = 4) Di-8-ANEPPS recordings from opposing ends of FDB fibers and the middle of the fiber. a) Example line scans from the same fiber on opposing ends. Red and green boxes show ROI’s for the fiber and the background, respectively. Yellow arrows mark shows the timing of stimulation. Even though the fiber exhibits an appreciable movement shortly after the stimulus, the sampled region of the line scan (within the red lines) remains within the fiber. b) Averages recordings from both ends and the middle of the same individual fibers. See text for fluorescence to voltage estimation. Yellow lines in the fiber cartoon show locations of line scan acquisition, approximately 50 µm from the ends of the fiber and the middle of the fiber. Note the lack of electrotonic potential in the middle recordings. Small movement artifacts can be seen after the end of the action potentials in these recordings. c) Averaged (n = 2) Di-8-ANEPPS recordings from the ends of the fibers in the presence of TTX. By blocking Na+ channels, we can measure the passive properties of the muscle cells.

### AP longitudinal propagation velocity calculated from the difference in AP time courses recorded at the same end of a skeletal muscle fiber for opposite polarity field stimulation

We have shown that we can optically obtain active responses from muscle fibers using field stimulation. With a dipole electrode, we stimulated Di-8-ANEPPS stained muscle fibers with alternating polarity. As noted earlier, this allowed us to initiate an AP from either end of the muscle fiber. Utilizing this, we set out to estimate the AP conduction velocity by recording from one end and switching the stimulation polarity.

We calculated the conduction velocity using the difference in t_1/2_ of the rising phases of the AP waveforms recorded at one end of the fiber during alternating polarity stimulation. By this method, we calculated the conduction velocity to be 0.39 ± 0.02 m/s. The average length of fibers was 446.2 µm. F_0_ during the first recording was compared to that of the final recording to determine the amount of bleaching (Figure S1). This difference in F_0_ was not found to be significant (Mann-Whitney, *p* = 0.94). Likewise, the peak ΔF/F_0_ of the first and last recordings on each polarity were compared to examine the signal rundown (Figure S2). Again, these differences were found to be not significant (Mann-Whitney, + Polarity: *p* = 0.56; - Polarity: *p* = 0.84). These results help show that the AP conduction velocity can be measured optically using Di-8-ANEPPS to record APs at a single end of a muscle fiber together with alternating polarity electrical field stimulation with bipolar electrodes.

### Calculating AP longitudinal propagation velocity using Ca^2+^ signals for alternating polarity field stimuli

Voltage recordings with Di-8-ANEPPS produce an observable electrotonic potential that will blend into the rising phase of the AP. Furthermore, the signals are relatively small, so signal averaging is needed to increase the signal to noise ratio. In contrast, using the low-affinity Ca2+ indicator mag-fluo-4 produces a clear, easily observable record of the calcium transient in a single (not averaged) record, with no stimulus artifact. As we have observed with the voltage signals, alternating polarity stimulation produces a calcium transient at one end which spreads to the opposing end due to the propagation of the AP depolarization [8]. Because of this, we can also use AP-evoked calcium transients to calculate the conduction velocity (Figure 4). A total of 25 fibers from 5 mice were used in this portion of the study. We calculated the conduction velocity using the difference in t_1/2_ of the rising phases of the AP-induced Ca^2+^ transients recorded at one end of the fiber during alternating polarity stimulation. The conduction velocity, when measured via calcium transients, is 0.37 ± 0.03 m/s. The average length of the fibers for these experiments was 436.6 µm. The difference in calculated conduction velocity between recordings performed using Di-8-ANEPPS and mag-fluo-4 (Figure 5) was not found to be statistically significant (Two sample t-test, *p* = 0.62). Thus, via remote bipolar electrodes Mag-fluo-4 can also be used to reliably measure the AP conduction velocity, and provides a much clearer signal.

**Figure 4.**
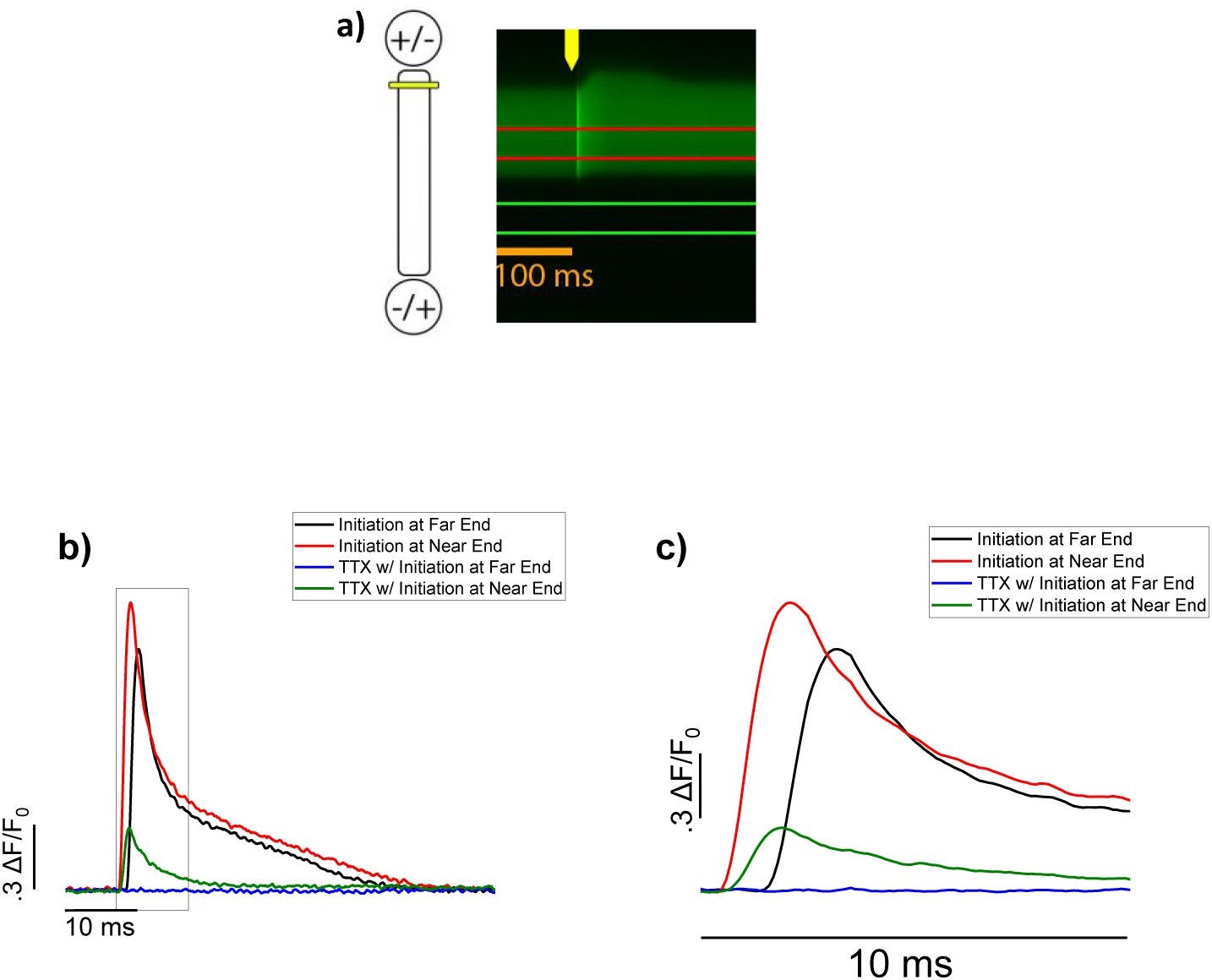
Example mag-fluo-4 recordings from one end of an FDB fiber. a) Yellow line shows relative location of line scan acquisition. Example line scan is shown with regions of interest for the fiber and the background (red and green, respectively). b) Resulting time course plots from line scans of one fiber. Recordings done with TTX show the local calcium release induced by passive depolarization, but not passive hyperpolarization. Significant differences were found between the peak amplitudes of the transients elicited by the positive and negative polarities (Paired sample T-test, + Polarity: x = 1.15; − Polarity: 1.26, N = 25, *p* = 0.03 × 10^-7^). c) Zoom-in of boxed in section of b) emphasizing the initiation of the calcium transient.

**Figure 5.**
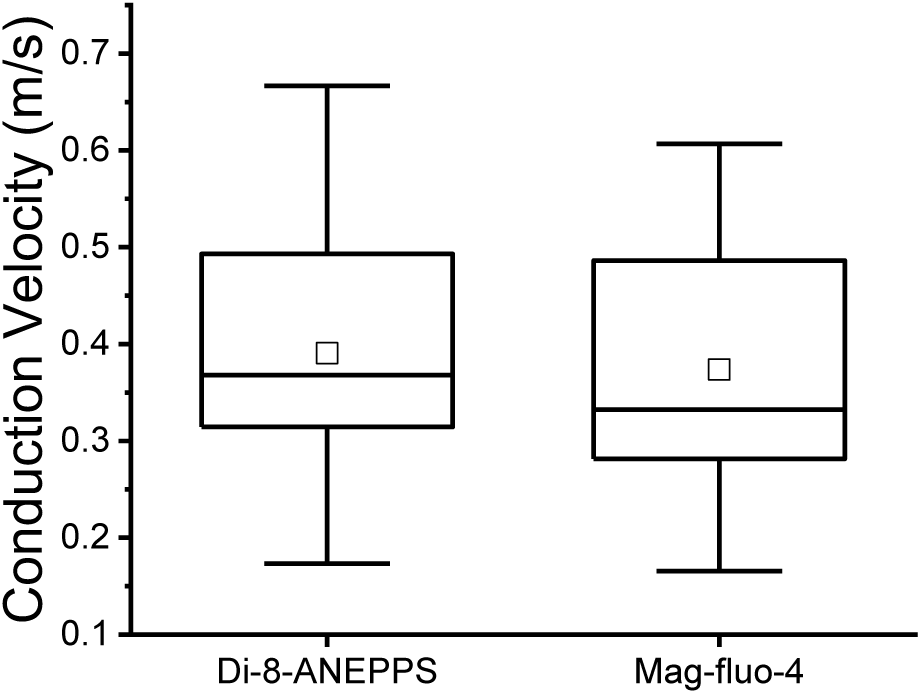
Conduction velocities measured by Di-8-ANEPPS and mag-fluo-4. Box represents Q1, median, and Q3. Whiskers show the minimum and maximum values. Smaller box displays the mean. No significant difference was found between the two conditions (Two sample T-test, *p* = 0.618).

### Conduction velocity is slowed in elevated extracellular potassium

Intramuscular increases in extracellular potassium concentrations have been seen in fatigue states [21]. Studies from other groups demonstrated that elevated extracellular potassium can depolarize the cell membrane due to a change in the Nernst potential for potassium [22, 23], hence altering sodium channel gating or slowing Na^+^-K^+^ pump activity [21, 24]. Due to these changes, we hypothesized that a slight increase of the extracellular potassium concentration could decrease the conduction velocity in skeletal muscle. Using mag-fluo-4, we tested the conduction velocity of each selected fiber first in our standard recording solution (L-15), which has a 5 mM concentration of KCl, then switched the media to a modified L-15 solution with 7.5 mM KCl, and tested at 5, 10, and 20 minutes after altering the KCl content (Figure 6a). At 5 minutes, we measured a significant decrease in conduction velocity in the elevated KCl group (n = 11) compared to control (One-way ANOVA, n = 15, *p* = 0.0003). This change persisted at 10 *(p* = 0.000002) and 20 minutes (*p* = 0.00000004, Figure 6b). Some of the cells became too depolarized to initiate Ca^2+^ transients at later time points, which accounts for the decrease in sample size as time goes on. Fibers exposed to a mock solution change (replacing the original 5 mM K solution) showed no change in conduction velocity at 5, 10 or 20 min (Figure 6b). These findings show that we can reliably track changes in conduction velocity using our methods.

**Figure 6.**
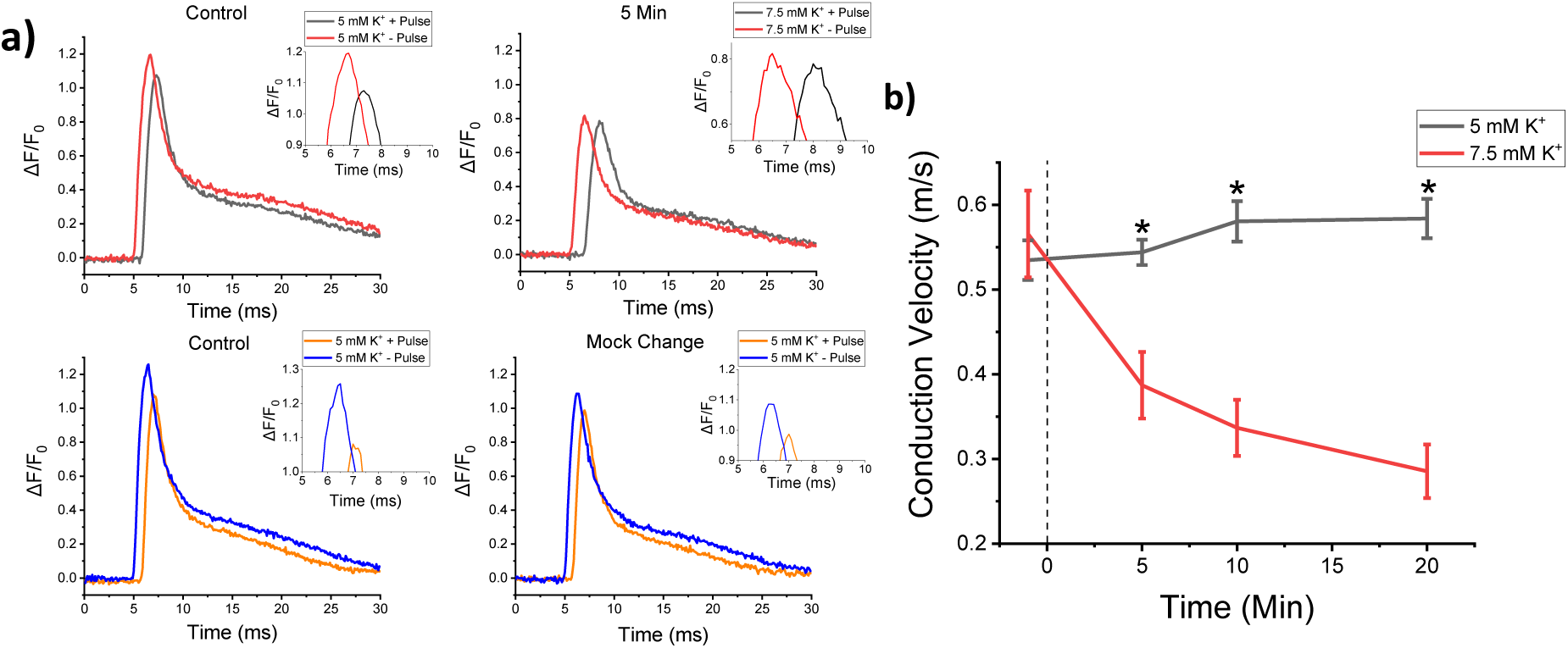
Conduction velocity measurements from WT fibers in 5 mM and 7.5 mM KCl using mag-fluo-4. a) Example Ca^2+^ transients from fibers in 5 mM and 7.5 mM KCl fibers. Inset shows a zoom-in the timing of the full-peak between transients recorded at the end of initiation versus the end of propagation. Fibers are shown before the solution switch and 5 minutes afterwards. b) Line & symbol plots show average changes in conduction velocity over time. Fibers that did not respond after an increase in KCl were removed. Dotted line indicates when solution was changed. Asterisk indicates significance established using a one-way ANOVA, 5 mM (n = 15) vs. 7.5 mK KCl (n = 11). Pre-Switch, *p* = 0.553; 5 min, *p* = 0.0003; 10 min, *p* = 0.000002; 20 min, *p* = 0.00000004).

### Modelling passive response of fiber to electric field stimulation by rem ote bipolar electrodes

Creating a model diagram for the passive electrical properties of the muscle fiber helps to understand and predict what would happen *in vivo* or *in vitro* during field stimulation via remote bipolar electrodes. We assume that in the immediate vicinity of a muscle fiber plated in the culture dish the following apply:

(1) The electric field generated in the bath by the remote bipolar stimulating electrodes is constant (i.e., the voltage changes linearly with distance along the direction of the field),

(2) The local electric field generated by the stimulating electrodes is locally spatially uniform in the vicinity of the fiber, and

(3) The presence of the fiber causes negligible change in the voltage generated in the bath by the stimulating electrodes.

Under these assumptions, the voltage outside the fiber will change linearly with distance along the fiber. If the fiber is exactly parallel to the field, the voltage gradient along the fiber will be the full voltage gradient parallel to the field. If the fiber is at an angle to the applied field, the voltage along the fiber will still change linearly with distance along the fiber, but voltage change per unit fiber length will be less than the full field by a constant geometric factor which depends on the fiber angle. In both cases we use the corresponding voltage applied to the low bath resistance to generate a linear voltage change along the outside of the fiber. We apply the extracellular voltage profile to a linear cable model of the fiber in order to calculate the fiber membrane potential as a function of time and location within the fiber.

Our analysis considers only fiber polarization due to the component of the applied electric field that is parallel to the long axis of the fiber. Fiber polarization due to the transverse component of the electric field is smaller because it only corresponds to a shorter distance (the fiber diameter) along the transverse field, in contrast to the fiber length which is subjected to the longitudinal component of the electric field. Transverse polarization is likely to be seen only in fibers oriented close to perpendicular to the applied field [8] or in fibers intentionally stimulated transversely using long electrodes parallel to the fiber.

### Passive cable model of an isolated skeletal muscle fiber in a large bathing solution

We represent the muscle fiber as a group of 7 longitudinal elements, each representing the membrane resistance and capacitance in 1/7 of the muscle fiber, flanked internally on each side by ½ the internal longitudinal resistance in 1/7 of the length of the fiber, and correspondingly flanked externally by the ½ of the external longitudinal resistance of the element (Fig. 7a), with the external longitudinal resistance set very low to maintain the constant field in the bath despite the presence of the fiber (condition 3, above). The values of the circuit elements were calculated from values of fiber internal resistance per unit length, membrane resistance times the unit length and membrane capacitance per unit length determined from papers describing electrical cable properties of mouse EDL skeletal muscle fibers [25]. Note that our cable model of the fiber ignores the effects of resistance in series with the T-tubules membrane capacitance, which we have not considered in our simulations to date.

**Figure 7.**
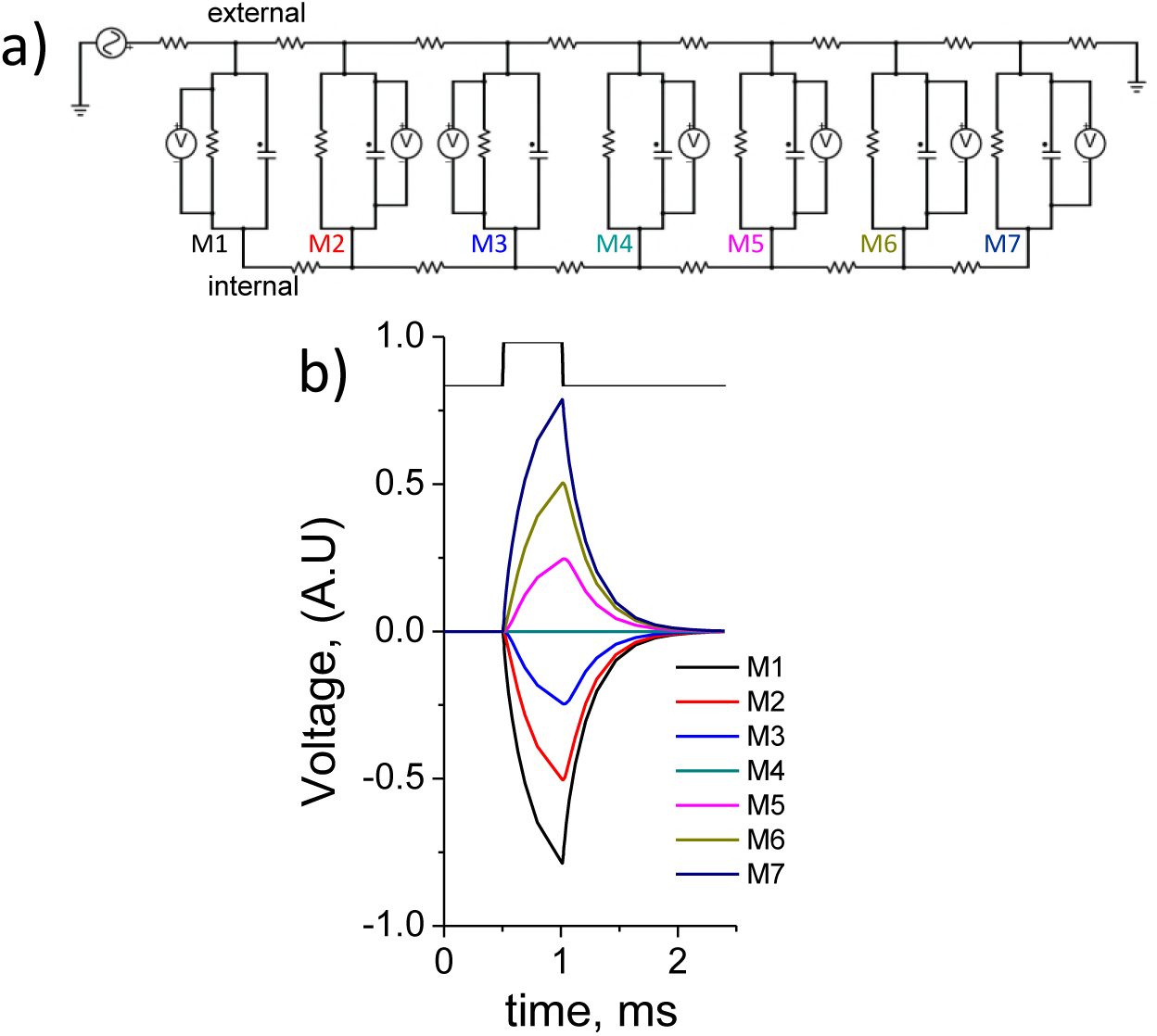
Circuit diagram for modeling the passive properties of FDB muscle fibers. a) Diagram was made via PartSim. Diagram was made to contain 7 identical elements. We have also constructed a model of a fiber 10 times longer. In the 10× model, the passive depolarization and hyperpolarizations have a larger amplitude closer to the ends, similar to this model, but there is a long segment in the middle where no changes in voltage are observed (not shown). The measuring points are color coded with the line graph in b, M1-M7; M4 is the element in the middle of the fiber. b) Example passive responses from the 7 element diagram. A 20 V pulse lasting 0.5 ms was applied to the circuit. The end of the circuit closer to the battery experiences depolarization, which degrades as the recording points move away from the battery. At the halfway point (M4-c), no voltage is recorded, but beyond that point a passive hyperpolarization is seen, and increases until the end of the circuit.

Simulations were done by applying a voltage step across the two end terminals of the extracellular series of resistances, and calculating the resulting voltage time course across each transmembrane element using the circuit solving software PartSim (AspenCore, LLC, 2017). For a 0.5 ms pulse applied to the bath via the field stimulating electrodes, the model exhibits a brief (0.5 ms) electronic depolarization at the fiber end nearest the negative electrode, and an equal but opposite polarity hyperpolarization at the end closest to the positive electrode (Fig. 7b). There is no electronic potential across the membrane elements at the fiber center, and successively decreasing polarization (depolarizing or hyperpolarizing) moving from the ends to the central elements, respectively. Thus the field stimulus produces depolarization over the half fiber facing the negative electrode, and hyperpolarization over the half facing the positive electrode, with maximum polarization at the fiber ends and none at the center. We observe similar depolarization and hyperpolarization at opposite ends of the muscle fibers *in vitro* using our system, indicating that our model is accurate. We have restricted the model to estimate the cable properties at 100 μm, but we are able to increase the length to test how these properties change with an increase in length. This model will help us quickly and reliably test the passive properties of different lengths and diameters of skeletal muscle fibers, allowing us to estimate what will happen to skeletal muscles *in vitro*.

## DISCUSSION

Isolated mouse muscle fibers are commonly used to assess various aspects of structure and function of muscle at cellular level. Physiological properties, signaling pathways, and more have been understood using this model [26–28]. Fluorescent probes and ultra-fast time lapse confocal microscopy approaches are among those that have been utilized to expand our knowledge of this system. These relatively non-invasive approaches have allowed us to more naturally observe and manipulate the endogenous systems. Our studies show that high-speed line scan potentiometric imaging allows us to examine, with high temporal resolution, the passive and active membrane voltage changes in response to field stimulation. Bipolar stimulation enables us to initiate APs at either end of the muscle fiber, which allows us to optically record the AP conduction velocity while recording only at a single end of the fiber. In addition, we are also able to examine conduction velocity with the calcium indicator mag-fluo-4. Previous research in our lab suggests that in the absence of AP initiation, the passive depolarization caused by bipolar field stimulation results in localized calcium release at the end of the fiber facing the cathode. This is dependent on the stimulation polarity, and results from a direct activation of the Ca_v_1.1 channels, the voltage sensors of the excitation-contraction coupling, at the depolarized end of the fiber and the subsequent non-propagated Ca2+ release [8]. Our current results support this finding. In our mag-fluo-4 recordings in the presence of TTX, a small amount of Ca2+ release can be seen (Figure 4), showing that passive depolarization indeed causes Ca2+ release.

Our model confirms some features of the electrotonic responses seen in the single muscle fibers. External field stimulation causes depolarization in the half fiber close to the negative electrode, and hyperpolarization over the half close to the positive electrode.

Conduction velocity of skeletal muscle is a useful physiological measure for examining how the intracellular and intermembrane elements could be functionally affected by different pathological conditions. This approach can be used to quickly and non-invasively obtain details about the muscle fiber excitability and AP properties. Previous studies have used microelectrodes and EMG recordings to examine conduction velocity [29–32]. Microelectrodes can measure membrane potential in a spatially restricted manner. For instance, measurements are limited to the surface membrane system and at the location of impalement. Moreover, while multiple electrodes could be used, these are still restricted to the surface membrane system. In addition, due to its invasive character, microelectrode techniques, especially with multiple impalements, may disrupt the normal cytoarchitecture and could alter normal responses. While EMG recording takes a global recording of an entire muscle group, it does not allow for the analysis of individual cellular components. Our system allows for the screening of individual muscle fibers’ AP properties occurring at both the surface and t-tubule system. The AP can be monitored simultaneously at multiple locations while allowing for high time and spatial resolution.

Optical recordings have their pitfalls as well. Estimating waveforms and conduction velocity from *xt* line scan images means that our temporal resolution is only as high as our line aquisition rate. In some microelectrode amplifiers the sampling rate can be similar or higher than in optical recordings. Additionally, the Di-8-ANEPPS signal itself is weaker than that of Ca^2+^ indicators, and while Di-8-ANEPPS is a potentiometric dye, it is not very voltage sensitive. A previous report showed that Di-8-ANEPPS only has a fluorescence change of 2.5% per 100 mV change [33]. In ordder to improve the signal to noise ratio, we took multiple recordings and averaged them (synchronized with the field stimulus pulse).

Mag-fluo-4 is much more sensitive to changes in calcium than Di-8-ANEPPS is to changes in voltage. Mag-fluo-4 provides a much clearer signal, without requiring averaging of multiple recordings. However, the propagation speed of Ca^2+^ transient is a derivate estimate of AP conduction. It is possible that as voltage travels along the membranes, the delay between the AP and Ca^2+^ transient could change. Therefore, in order for these measurements to maintain validity, we must assume that the delay between APs and Ca^2+^ transients is the same at both ends of the fiber.

Conduction velocity analyses in frog muscle have shown much higher speeds than what we observed in our studies. A classic study using winter and spring frogs found average conduction velocities of 2.44 m/s and 2.05 m/s, respectively [34]. Conduction velocity is known to increase with fiber diameter and length, and the average frog muscle is much larger than the mouse muscles we observed in our studies. Studies in rat have also shown higher values that what we recorded, perhaps for similar reasons. A study from Kupa and colleagues calculated the conduction velocity in rat EDL fibers to be 3.02 m/s on avarage, and 1.70 m/s on average in the soleus [32].

The AP conduction velocity of 0.39 m/s obtained here in mouse FDB muscle fibers is about an order of magnitude lower than the values 3.5 m/s and 3.8 m/s previously reported for mouse Sartorius [35] and EDL [36] muscle fibers. A key difference between the fibers studied here and those described in earlier reports is that FDB fibers are much shorter than the previously studied fibers. In a much shorter fiber, a proportionally slower AP can produce a similarly synchronized depolarization at the fiber ends as produced by a proportionally faster AP in a much longer fiber. Thus, slower conduction is sufficient for near synchronous activation of the relatively short FDB fibers used here. Furthermore, a reduced conduction velocity due to a decreased level of membrane Na^+^ channels would decrease Na^+^ influx, which spares energy needed to reverse the Na^+^ influx via the Na/K pump. We thus hypothesize that short muscle fibers may be specialized to use slower propagating APs for energy conservation. A lower Na^+^ channel density in FDB fibers could also explain their relatively high sensitivity to K^+^ depolarization. We have observed a decreased conduction velocity to about 49% of control and some failure of excitation for the modest increase of extracellular [K+] from 5 to 7.5 mM in FDB fibers (Fig. 6). Furthermore, we found that 10 mM extracellular [K+] completely eliminated excitability of FDB fibers (not shown), which was not observed in reports on longer muscle fibers from mice [36]. The increased susceptibility of FDB fibers to suppression of excitability by elevated extracellular K^+^ would be consistent with a lower level of Na^+^ channel membrane expression in FDB fibers compared to longer muscle fibers.

Previous studies that have examined sarcolemmal conduction velocity (ie, the longitudinal conduction velocity along the muscle fibers) in humans found much higher values, up to 6.4 m/s [36, 37]. As noted in the introduction, the radial conduction velocity in the T-tubules is much slower. In *Xenopus,* this value has been measured at <0.3 m/s [38]. With respect to the z-axis, our recordings were taken from the middle of the fiber, suggesting that we were recording the T-tubule membrane potential, and thus the longitudinal speed of spread of T-tubule depolarization along the fiber. Granted, all of these studies were conducted in different species, but these provided a good reference point for our studies.

Our recordings with both Di-8-ANEPPS and mag-fluo-4 provided similar values for the longitudinal conduction velocity. The membranes of the T-tubule and the sarcoplasmic reticulum (SR) are molecularly coupled [39], and T-tubule membrane depolarization leads to activation of the SR calcium release channels [40]. SR calcium release cannot occur without T-tubule voltage propagation. Because of the small time window between depolarization, calcium release and the mag-fluo-4 signal, and by extension Ca2+, we can also use the mag-fluo-4 siganl to estimate the AP conduction velocity. In our testing conditions, we observed that both mag-fluo-4 and Di-8-ANEPPS are useful tools to measure AP conduction velocity. While we did not observe any significant differences between Ca^2+^ and voltage propagation, we cannot guarantee this will be the case in every experimental condition.

When field stimulating fibers using remote bipolar electrodes as under these conditions we have to take the angle of the fiber relative to the electric field into account. In our diagrams (Figure 1), the fiber is oriented parallel to the line between the 2 electrodes. When in this orientation, the fiber experiences the largest electrotonic electrical polarization during field stimulation. As the fiber rotates away from parallel, the amplitude of the electronic potentials, decreases until the fiber is perpendicular to the electric field, where it is least responsive due to zero electrotonic potential at the fiber ends but with a small transverse polarization due to the field which is now across the fiber. Using different types of electrode geometry will generate different electric fields, eliciting different responses. If we use a focal electrode, we can elicit depolarization from any point along the fiber, from which an AP would propagate in both directions [8].

A previous paper from our lab characterized the behavior of fibers that only contract and display Ca^2+^ transients at one end when stimulated with a given polarity [8]. If a fiber only twitches at one end, it cannot be used for conduction velocity measurements using mag-fluo-4. There will only be local calcium release at the twitch site, and no local or propagated AP. However, alternating fibers can still be used to examine the passive electrical properties of the fiber using Di-8-ANEPPS. The electrotonic potential will still be produced in alternating fibers. Whether using alternating fibers or uniformly contracting fibers, our approaches can show the passive and/or active properties of skeletal muscle fibers, using either mag-fluo-4 or Di-8-ANEPPS.

On the whole, this approach allows us to attain good signal to noise ratios using Di-8-ANEPPS. This approach can also be applied to other cell types, such as cardiac or axonal projections. There are some relatively new voltage or calcium-sensitive sensors that could work very well in tandem with our method. Genetically engineered fluorescent probes have been shown to be useful and efficient in measuring membrane potential and calcium changes. ASAP1, a fast voltage sensor [41] and G-CaMP, a genetically encoded calcium probe [42], has been shown to be effective in neurons [43], however the time response of Di-8-ANEPPS (<1ms) is superior to that of genetically encoded voltage or calcium probes. Yet, due to dye uptake and delocalization, Di-8-ANEPPS is not suitable for long-term experiments (>4 hours).

In our experiments with fibers challenged with extracellular elevated K^+^, we observed that conduction velocity slowed in a time dependent manner. In fatigue states, where extracellular K^+^ is elevated, the cell membrane is partially depolarized, the sodium permeability is reduced, and the ability to generate APs is decreased. We suspect that something similar is happening in our conditions as well, despite the lack of K^+^ efflux from the intracellular milieu. KCl is a known membrane depolarizer, and the depolarization effect should occur immediately. Furthermore, a notable percentage of the fibers became unable to generate APs, even at just 7.5 mM KCl. This percentage also increased with time. Regardless, these changes demonstrate that we can track alterations in conduction velocity using fluorescent dyes coupled with field stimulation.

## CONCLUSIONS

We have demonstrated that Di-8-ANEPPS and mag-fluo-4 can be used to measure the AP conduction velocity using high speed optical recording at one end of the fiber along with alternating polarity field stimulation from remote bipolar electrodes. Mag-fluo-4 provides much clearer signals, so providing the muscle fiber is producing calcium transients, this is the easier and more reliable indicator to use. We will be able to use this technique to examine the effect of different diseases and cellular states on the AP propagation. For example, how does T-tubule dysregulation or dysfunction affect the propagation? If a membrane protein is altered in diseases models, how does that affect the propagation? We anticipate that this novel but simple method will be an important additional research tool to reexamine neurological diseases such as amyotrophic lateral sclerosis, Alzheimer’s and Huntington’s. These pathological conditions were considered to arise from or to primarily affect the nervous system, however, numerous reports have shown that skeletal muscle is also implicated in the pathophysiology of these diseases [44–46].

Research reported in this publication was supported by the National Institute of Arthritis and Musculoskeletal and Skin Diseases of the National Institutes of Health under Award Number R37-AR055099 (to M. F. S.). Q.B. was supported by NIAMS-NIH training grant T32 AR007592 to the Interdisciplinary Program in Muscle Biology, University of Maryland School of Medicine. The content is solely the responsibility of the authors and does not necessarily represent the official views of the National Institutes of Health.

## Author Contributions

Q.B.: Experimental design, performed the majority of experiments and analysis; drafted and approved the manuscript

S.J.P.: Performed initial experiments with calcium indicator, developed methods and approved the manuscript

S.R.I: Developed methods, edited and approved the manuscript R.M.L.: Developed methods, edited and approved the manuscript

E.O.H.: Experimental design and approach, developed circuit model, edited and approved the manuscript

M.F.S.: Conceived the project, experimental design and approach, developed circuit model, edited and approved the manuscript

---

**Figure S1.**
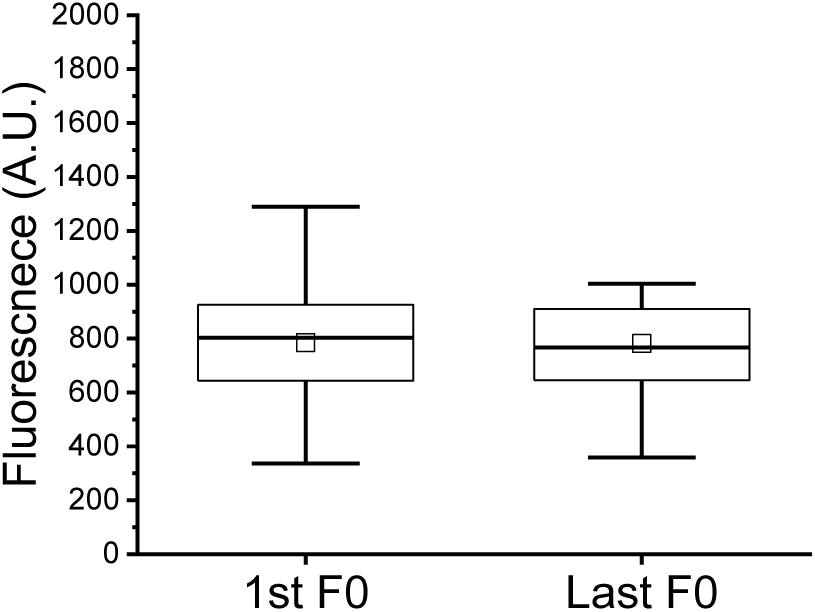
Box chart displaying data from bleaching examination in Di-8-ANEPPS. Boxes show Q1, median, and Q3. Smaller box represents the mean. Whiskers show the range of values. No significant difference between the 1st stimulus and final stimulus was found (Mann-Whitney, *p*= 0.935).

**Figure S2.**
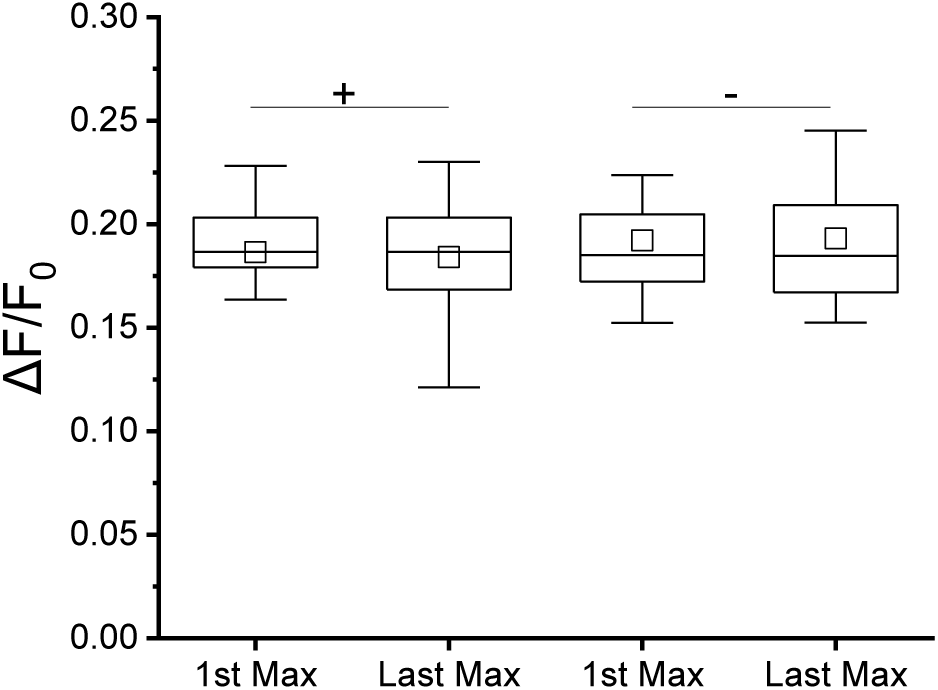
Box charts displaying signal rundown between conditions when recording using Di-8-ANEPPS. Boxes show Q1, median, and Q3. Smaller box represents the mean. Whiskers show the range of values. No significant difference was found between the first and last stimulation of either polarity (Mann-Whitney: + Polarity: *p* = 0.57; − Polarity: *p* = 0.84).

**Table S1.** Parameters recorded from isolated WT FDB fibers using Di-8-ANEPPS or mag-fluo-4. Parameters are expressed as mean ± SEM where appropriate. Differences between recordings of conduction velocity using Di-8-ANEPPS and mag-fluo-4 were not significantly different (Two Sample T-test, *p* = 0.618).

| Parameter | Di-8-ANEPPS Recordings | Mag-fluo-4 Recordings |
| --- | --- | --- |
| Time to Half Peak (+ Polarity) | $1.8 \pm 0.29$ ms | $11.3 \pm 0.08$ ms |
| Time to Half Peak (– Polarity) | $0.89 \pm 0.29$ ms | $10.3 \pm 0.09$ ms |
| Conduction Velocity with Half Peak | $0.390 \pm .02$ m/s | $0.373 \pm 0.025$ m/s |
| Average Fiber Length | $347 \pm 11$ μm | $437 \pm 16$ μm |
| n, m | 30, 12 | 25, 5 |

**Table S2.** Conduction velocities of experimental FDB fibers as recorded with mag-fluo-4 using the control external solution (5 mM K+) or at different times after challenging the fibers with a test solution with 7.5 mM K+. Conduction velocities are represented as mean ± SEM, n, m.

| Condition | Measured by mag-fluo-4 |
| --- | --- |
| 5 mM K<sup>+</sup> FDB | $0.566 \pm 0.05$, 10, 3 |
| 7.5 mM K<sup>+</sup> FDB – 5 min | $0.387 \pm 0.04$, 10, 3 |
| 7.5 mM K<sup>+</sup> FDB – 10 min | $0.337 \pm 0.03$, 10, 3 |
| 7.5 mM K<sup>+</sup> FDB – 20 min | $0.285 \pm 0.03$, 10, 3 |

